# Differential Survival of Retinal Ganglion Cell Types Following Branch Retinal Vein Occlusion

**DOI:** 10.64898/2026.10.20.760463

**Authors:** Saman Behboudi Tanourlouee, Chao Ren, Andrea Santeford, Sean McCracken, Minglei Zhao, Christopher Zhao, Rajendra S. Apte, Chao Zhou, Philip R. Williams

## Abstract

**Purpose:** Branch retinal vein occlusion (BRVO) is a common cause of retinal ischemia, rapidly inducing neuronal death and often producing irreversible vison loss. Interventions such as thrombolysis restore perfusion but cannot recover neurons already lost, and most patients retain permanent visual deficits. Identifying which neuronal populations die, and how rapidly, may therefore reveal timing windows and targets for neuroprotective treatment. To determine the timing of pathology, we evaluated vascular changes and neuronal degeneration in a photocoagulation-induced mouse model of BRVO. To determine differences in retinal ganglion cells (RGC) type susceptibility, we tracked the survival of RGC subpopulations using immunofluorescence markers.

**Methods:** BRVO was induced by Rose Bengal injection followed by targeted photocoagulation of a branch vein. Obstruction was confirmed by fluorescein angiography and vascular damage assessed by optical coherence tomography angiography. Cell survival was quantified by immunostaining at 3, 7, and 14 days using RNA-Binding Protein with Multiple Splicing (RBPMS, pan-RGC). RGC subtype markers for intrinsically photosensitive RGCs (TBR2), α-RGCs (SPP1), ON–OFF direction-selective RGCs (CART), and F-RGCs (FOXP2) were used to compare subtype survival.

**Results:** Occlusion selectively disrupted the superficial and intermediate capillary plexus, sparing the deep plexuses. All examined neuronal populations were significantly depleted, but with markedly different magnitude and timing. α-RGCs and F-RGCs were the most vulnerable subtypes, both approaching complete loss by one week, whereas intrinsically photosensitive RGCs were the most resistant.

**Conclusions:** α-RGCs, typically resilient in glaucoma, trauma, and excitotoxicity, were markedly vulnerable in BRVO, indicating that RGC subtype susceptibility is insult-specific.

## 1. Introduction

The retina is one of the most metabolically active tissues in the body, with one of the highest oxygen consumption rates per gram, exceeding that of the brain.^1,2^ Thus, the retina is extremely sensitive to disruption of blood circulation. Many leading causes of irreversible vision loss, including diabetic retinopathy, glaucoma, and age-related macular degeneration, involve vascular dysfunction, impaired retinal oxygen homeostasis, and hypoxia-driven neurovascular degeneration.^3–6^ Anatomically, retinal ischemia exists on multiple scales from capillary ischemia to complete retinal ischemia. Branch retinal vein occlusion (BRVO) is a regional form of retinal ischemia, the predominant subtype of retinal vein occlusion, and the second most common retinal vascular disorder after diabetic retinopathy.^7,8^

BRVO usually starts with compression of the retinal vein at arteriovenous crossings due to retinal artery thickening and arteriosclerosis, which puts mechanical pressure on the vein. Another etiology is venous thrombosis, which can occur in patients with hypercoagulable states or coagulation disorders.^9^ The resulting increase in venous pressure acutely causes hemorrhage, capillary leakage, and macular edema. Persistent venous congestion after BRVO impairs retinal perfusion and leads to retinal hypoxia, which activates HIF-1a and increases VEGF expression. Increased VEGF can promote vascular permeability and neovascularization, while elevated venous pressure and inflammation further contribute to blood–retinal barrier disruption and retinal edema.^10–13^ Although BRVO onset is acute, visual dysfunction evolves subacutely. Retinal edema involving the macula causes an early decrease in visual acuity, which can improve over weeks to months depending on BRVO subtype.^9,14^ Unlike arterial occlusive diseases, BRVO lacks a clinically established acute therapeutic window targeting the initial ischemic insult. Instead, treatment has historically been guided by disease progression, with the Branch Vein Occlusion Study initially supporting a 3-month observation period before laser intervention for persistent macular edema.^15^ Management has shifted from grid laser photocoagulation to intravitreal therapies, including anti-VEGF agents, as well as corticosteroids, which target vascular leakage and macular edema.^16–19^ Anti-VEGF agents are also used to treat proliferative neovascularization in conjunction with targeted sectoral retinal photocoagulation to reduce the ischemic retinal burden. There are currently no available therapies that selectively target or reverse neuronal degeneration and relatedvisual field deficits associated with BRVO.

Despite extensive characterization of vascular changes and hemodynamics in BRVO, comparatively little is known about how individual neuronal subtypes within the inner retina respond to and survive venous-occlusive hypoxia. Retinal ganglion cells (RGCs) are comprised of many molecularly and functionally distinct subtypes with variable metabolic demands, dendritic architectures, and susceptibility to degeneration in glaucoma and optic neuropathy models.^20–23^ Importantly, the clinical timing framework for BRVO is derived primarily from vascular and edema resolution, reflecting the stage at which retinal nonperfusion is clinically recognized rather than directly identifying when neuronal injury becomes irreversible.^24^ Current clinical endpoints, including macular edema and visual acuity, reflect the overall consequences of retinal vascular occlusion but do not directly distinguish vascular pathology from neuronal injury. Consequently, the contribution and temporal dynamics of neuronal damage remain difficult to define clinically.^25, 26^ If neuronal populations have different timing of cell loss relative to edema and vascular defects after ischemic injury, the window for neuroprotective treatment may not be the same as the current treatment window for vascular damage in BRVO. Additionally, neuronal injury may continue even after the initial ischemic event, which could contribute to persistent visual deficits despite successful anatomical resolution of macular edema.^27^ Understanding the timing of cell loss may therefore be important for identifying when neuroprotective treatment could be most effective in BRVO. Different RGC populations may also have different levels of vulnerability to injury. In glaucoma, different RGC subtypes show different levels of vulnerability to injury. A mechanistic understanding of these differences has led to the development of subtype-specific biomarkers and neuroprotective strategies that function in animal models.^28–30^ Whether a similar pattern of cell-type-specific vulnerability occurs in BRVO remains unknown. Here, we investigate patterns of RGC vulnerability following BRVO to define the types most affected by ischemia.

## 2. Methods

### 2.1 Animals and Induction of BRVO by Rose Bengal

Experiments were performed on C57BL/6J (Jackson Laboratory 000664) and Cx3cr1-GFP (Jackson Laboratory 005582) mice of both sexes, aged 8–12 weeks. Animals were randomly assigned to an ischemia group (photocoagulation-induced branch retinal vein occlusion, BRVO) or a sham group. All procedures were approved by the Institutional Animal Care and Use Committee (IACUC) and conducted in accordance with the ARVO Statement for the Use of Animals in Ophthalmic and Vision Research. Mice were anesthetized with an intraperitoneal injection of ketamine/xylazine (60 mg/kg and 10 mg/kg, respectively), and pupils were dilated with topical atropine sulfate (1% w/v). To enable simultaneous vessel visualization and photothrombotic occlusion, animals in the ischemia group received a retro-orbital injection (150 µL per eye) of a mixed solution of FITC-dextran (10 mg/mL in PBS; MW 2,000 kDa, Sigma Aldrich # FD2000S) and Rose Bengal (8 mg/mL in PBS; Thermo Scientific #189450250). Sham animals received retro-orbital FITC-dextran alone, without Rose Bengal, and were otherwise handled identically. A single retinal branch vein was targeted for photothrombosis under an epifluorescence microscope (Olympus/Zeiss [specify model]) equipped with a Mitutoyo 20× long-working-distance objective and a Texas Red filter set to excite Rose Bengal. The target vein was illuminated at a power of 1 mW out of the microscope objective for 18 minutes to induce localized endothelial photo-oxidative injury and thrombotic occlusion. Body temperature was maintained at 37°C throughout the procedure using a heating pad, and corneal hydration was preserved with ophthalmic lubricant (GenTeal gel), which also corrected for optical aberration of the front of the eye.^31^ Successful occlusion was confirmed intraoperatively by cessation of blood flow and post-thrombotic vessel dilation/tortuosity proximal to the occlusion site under fluorescence observation and subsequently verified by fluorescein angiography (see below).

### 2.2 In vivo retinal imaging

To confirm occlusion and monitor vascular changes longitudinally, fluorescein angiography (FA), and optical coherence tomography angiography (OCTA) were performed on all animals immediately before photocoagulation (baseline) and at 2 days post-occlusion. FA was acquired using a Micron III retinal imaging system (Phoenix-Micron, Inc.) following retro-orbital FITC-dextran injection, to characterize the caliber and course of the occluded branch vein and adjacent arterioles. The OCTA imaging system is a custom SD-OCT system, at 850±80 nm, with an axial resolution of ~2.3 µm in tissue, and a sensitivity of ~97 dB, respectively.^32^ For capillary layers, we binarized the vessel maps from OCT-A *en face* projection and calculated the vessel density index (VDI) was calculated as the ratio of area occupied by vessels. The local perfusion density map was then generated by defining a sliding window kernel and convolving it with the binary vessel image to compute the local fraction of vessel pixels. The fractal dimension was calculated using a box-counting method. This process involves defining logarithmically spaced box sizes as powers of two, sliding these boxes across the binary image, and tallying the number of boxes that contain at least one vessel pixel. Finally, linear regression was used to determine the slope of the log-log relationship between the box counts and the inverse of the box sizes, which yields the final fractal dimension.

### 2.3 Tissue collection and immunohistochemistry

At 3, 7, and 14 days post occlusion (DPO), mice were deeply anesthetized by intraperitoneal injection of tribromoethanol (Avertin; 250mg/kg; Sigma-Aldrich, T48402) and transcardially perfused with ice-cold phosphate-buffered saline (PBS) followed by 100 mL of 4% paraformaldehyde (Sigma-Aldrich) in PBS. Eyes were enucleated and post-fixed by immersion in 4% PFA in PBS at 4°C for 30 minutes. Following dissection of the anterior segment and lens, whole retinas were isolated and stored in PBS at 4°C until further processing. For cryoprotection, retinas were equilibrated overnight in 30% sucrose in PBS at 4°C and subjected to three freeze–thaw cycles on dry ice against a glass slide to enhance antibody penetration. Retinas were then washed three times for 10 minutes in PBS at room temperature and blocked for 1–3 hours at room temperature in blocking solution consisting of 10% normal horse serum (Sigma-Aldrich, S-2888) and 0.5% Triton X-100 (Sigma-Aldrich, 11332481001) in PBS. Retinas were incubated in primary antibodies diluted in blocking solution for 5–7 days at 4°C on an orbital shaker. The primary antibodies used in this study are listed in Table 1.

**Table 1.**
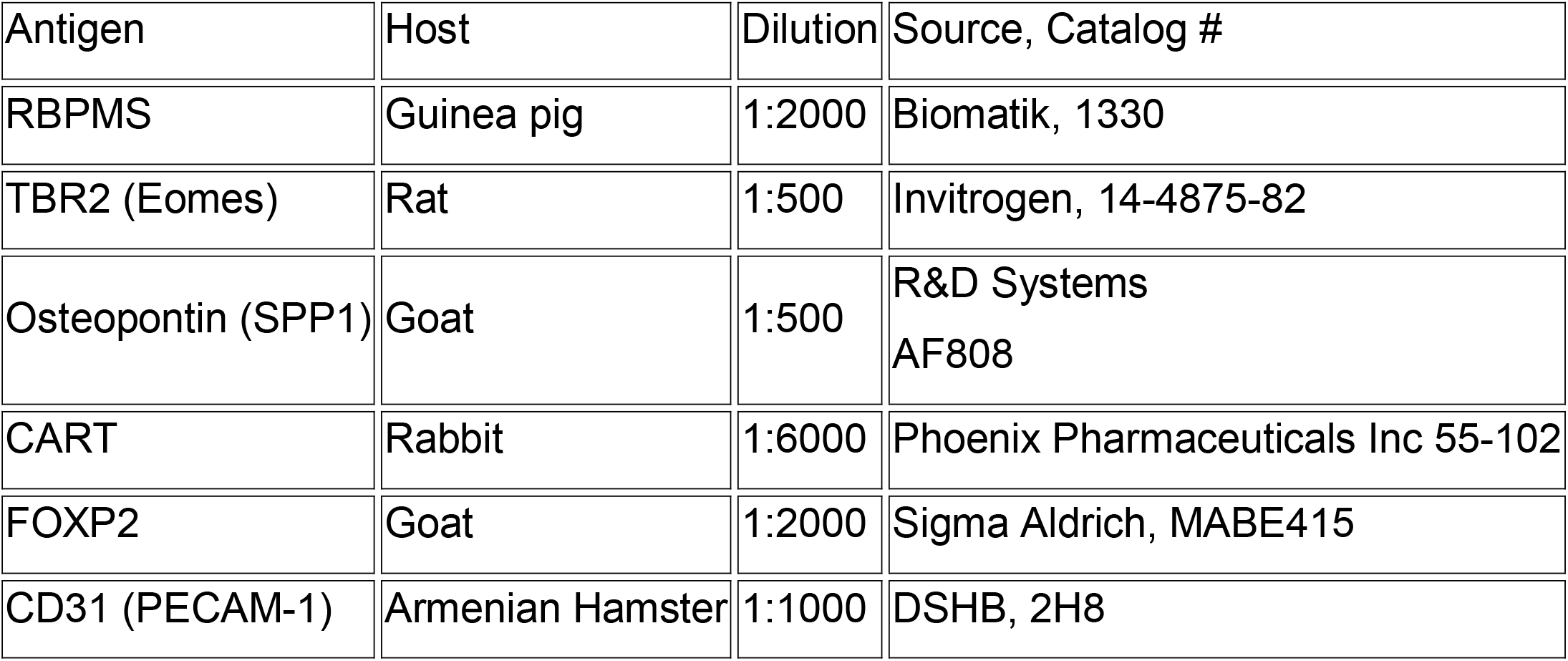
Description of the primary antibodies used in this study.

Following primary antibody incubation, retinas were washed three times for 10 minutes each in PBS at room temperature and incubated for 2–3 days at 4°C in species-appropriate secondary antibodies (donkey anti-guinea pig/rat/goat/rabbit, raised against the corresponding primary host species, cross-adsorbed, and conjugated to Alexa Fluor 405, 488, 568, or 647; Jackson ImmunoResearch), each diluted 1:1000 in blocking solution. For some samples, DAPI (Sigma Aldrich, MBD0015) was added to the secondary antibody solution at a dilution of 1:10,000. Retinas were washed three times for 10 minutes each in PBS, flat-mounted onto glass slides (ganglion cell layer up), coverslipped with Vectashield Antifade Mounting Medium (Vector Laboratories, H-1000-10), and sealed with nail polish.

### 2.4 Confocal imaging

Whole-mounted retinas were imaged on a Zeiss LSM 710 laser-scanning confocal microscope equipped with a 20× air objective (Zeiss Plan-Apochromat, 0.8 NA). Image stacks were acquired at 3.5 μm z-spacing (35-50 μm total z stack depth) and 0.446 μm xy pixel resolution. For scanning vasculature and microglia, we used higher resolution (0.32 μm xy pixel). A 6 × 6 tiled montage centered on the optic nerve head was collected for each retina, yielding a total imaged field of 3150 μm × 3150 μm, and tiles were automatically stitched using Zeiss ZEN Blue software. For retinal thickness measurements, two regions measuring 319.45 × 319.45 μm each were scanned per retina, with a z-step interval of 0.43 μm (90-130 μm total z stack depth).

### 2.5 Image analysis and cell quantification

Cell segmentation and quantification were performed on 0.5 mm^2^ retinal regions peripheral to the photocoagulation site, directly contralateral to the site for the non-occluded region control, or at a similar centro-peripheral location in sham samples. Z-stack images were batch-processed automatically with a custom Python pipeline that generated mean-intensity projections, segmented RBPMS^+^ cells with Cellpose (cpdino model), and exported the outlines as Fiji ROIs. After manual removal of false-positive ROIs, marker intensity was measured in each ROI after local background subtraction. Five clearly positive RBPMS^+^ cells per marker and image set the positivity threshold, single- and double-positive cells were counted as a percentage of RBPMS^+^ cells, and all classifications were manually checked; the scripts are available at [<u>GitHub repository</u>].

## 3. Results

### 3.1 Local capillary destruction and inflammatory response after BRVO

We used fluorescein angiography to monitor retinal artery and vein branches before and after photocoagulation to validate the vein occlusion. Two days after BRVO, the targeted vein showed complete occlusion, while other branch arteries and veins showed no signs of occlusion (Fig. 1A upper). We OCTA (n=3 per group) at 2 DPO (Fig. 1A lower) To evaluate retinal capillary bed perfusion in superficial, intermediate and deep vascular plexuses (SVP, IVP, DVP). OCTA imaging demonstrated complete elimination of blood flow within the occluded branch vein together with destruction of the surrounding capillary network (Fig. 1B). Capillary loss was most prominent within the superficial and intermediate capillary plexus in the affected retinal region, whereas the remaining retinal vasculature retained its normal morphology and vessel density (Fig. 1 B).

**Figure 1.**
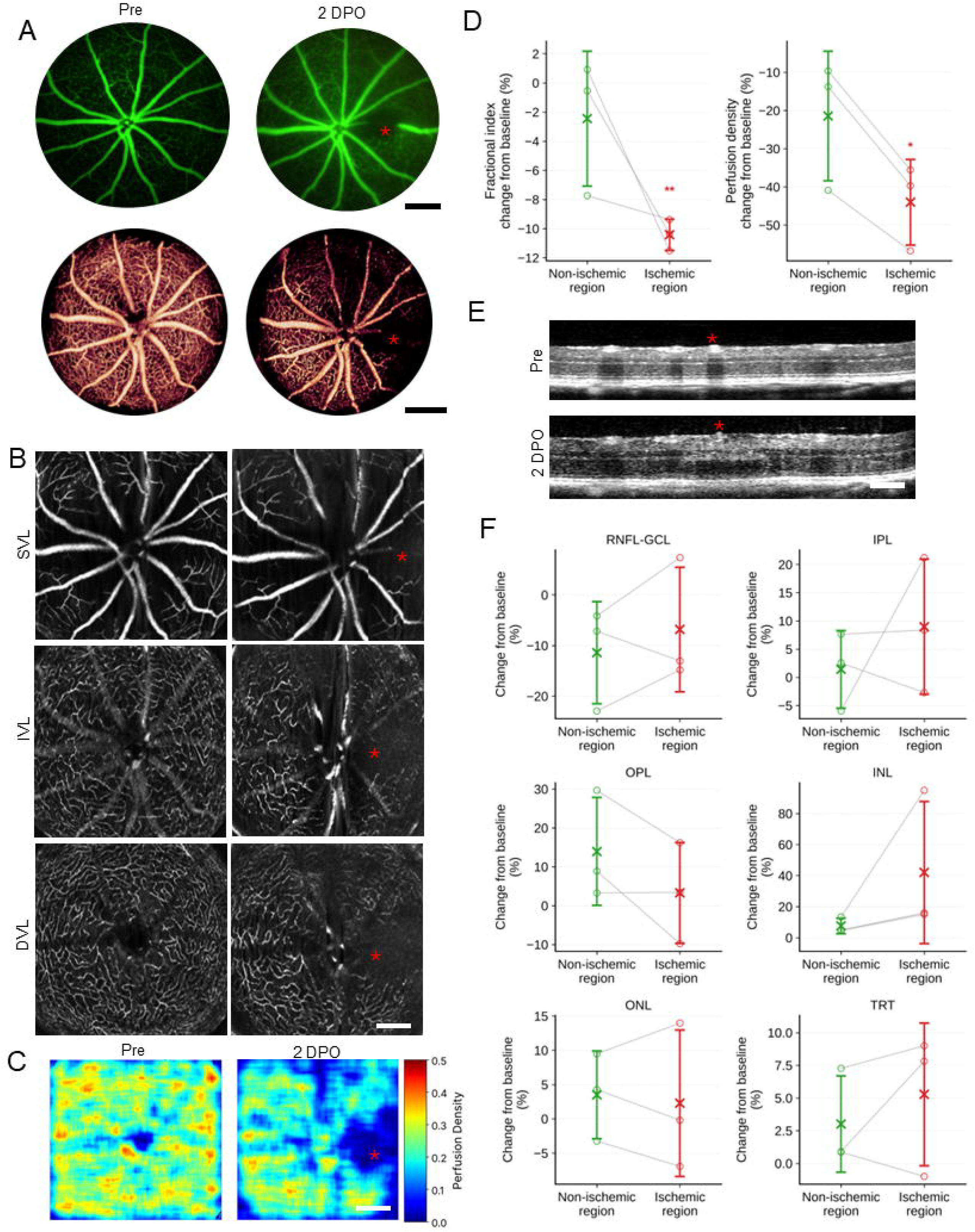
Retinal angiography and OCTA changes following photocoagulation-induced BRVO. (A) Retinal fluorescein angiography and OCTA (scale bars = 300 µm) before and 2 days after photocoagulation-induced BRVO, showing complete vein occlusion (red asterisks throughout). (B) OCTA scans of SVL, IVL, and DVL vascular plexuses before and after BRVO (Pre and 2 DPO). Scale bar = 200 µm. (C) Retinal perfusion maps before and after BRVO. Scale bar = 300 µm. (D) Quantification of retinal fractional index (left), and perfusion density (right) changes in the occluded and non-occluded regions. (E) OCT cross-sectional images of retinal layers before and 2 DPO after photocoagulation. Scale bar = 180 µm. (F) Quantification of changes from baseline in retinal sublayer and total retinal thickness in occluded and non-occluded regions over 2 days following BRVO.

To compare blood flow dynamics between occluded and non-occluded regions of the retina, we divided the retina into equal hemispheres and compared pre versus post perfusion density and vascular fracture for each hemisphere. Perfusion density and vascular fracture dropped, almost 40% and 10% respectively (p=0.021, p=0.0035, n=3 per group) at 2 days post coagulation in occluded regions. BRVO also caused a non-statistically significant hypoperfusion and decrease in vascular fraction in the non-occluded hemisphere. Comparison between the two hemispheres showed perfusion density was significantly lower in occluded areas compare to contralateral region, but vascular fracture was not statistically different (Fig. 1 C, D).

To examine changes in neural retina, cross-sectional OCT images were analyzed and confirmed complete absence of perfusion within the targeted vein (Fig. 1E). Although none of the changes were statistically significant, we observed evidence of edema in that total retinal thickness (TRT) increased slightly in both regions: +5.3 ± 5.5% (mean ± stdev) in the ischemic region and +3.0 ± 3.7% in the non-ischemic region (n = 3). (Fig. 1F). In contrast, the nerve fiber layer-ganglion cell layer complex (RNFL-GCL) became thinner in both regions: −6.8 ± 12.3% in the ischemic region and −11.4 ± 10.1% in the non-ischemic region (Fig. 1F), suggesting early onset RGC death.

To examine vasculature changes after BRVO with higher spatial and temporal resolution, we immunostained retinal wholemounts with the endothelial cell marker CD31 at 3, 7 and 14 DPO (Fig. 2). In contrast to OCTA measurements, at 3 DPO the occluded regions showed no obvious changes in vascular density or morphology (Fig. 2B). However, by 7 DPO, marked destruction of the capillary bed surrounding the occluded vessels was observed, which subsequently resulted in a localized reduction in capillary density within the occluded regions at 14 DPO (Fig. 2C–D). Importantly, we did not identify any vascular abnormalities in the non-occluded regions of BRVO retinas (Fig. 2A), indicating that our photocoagulation treatment was constrained to the targeted area.

**Figure 2.**
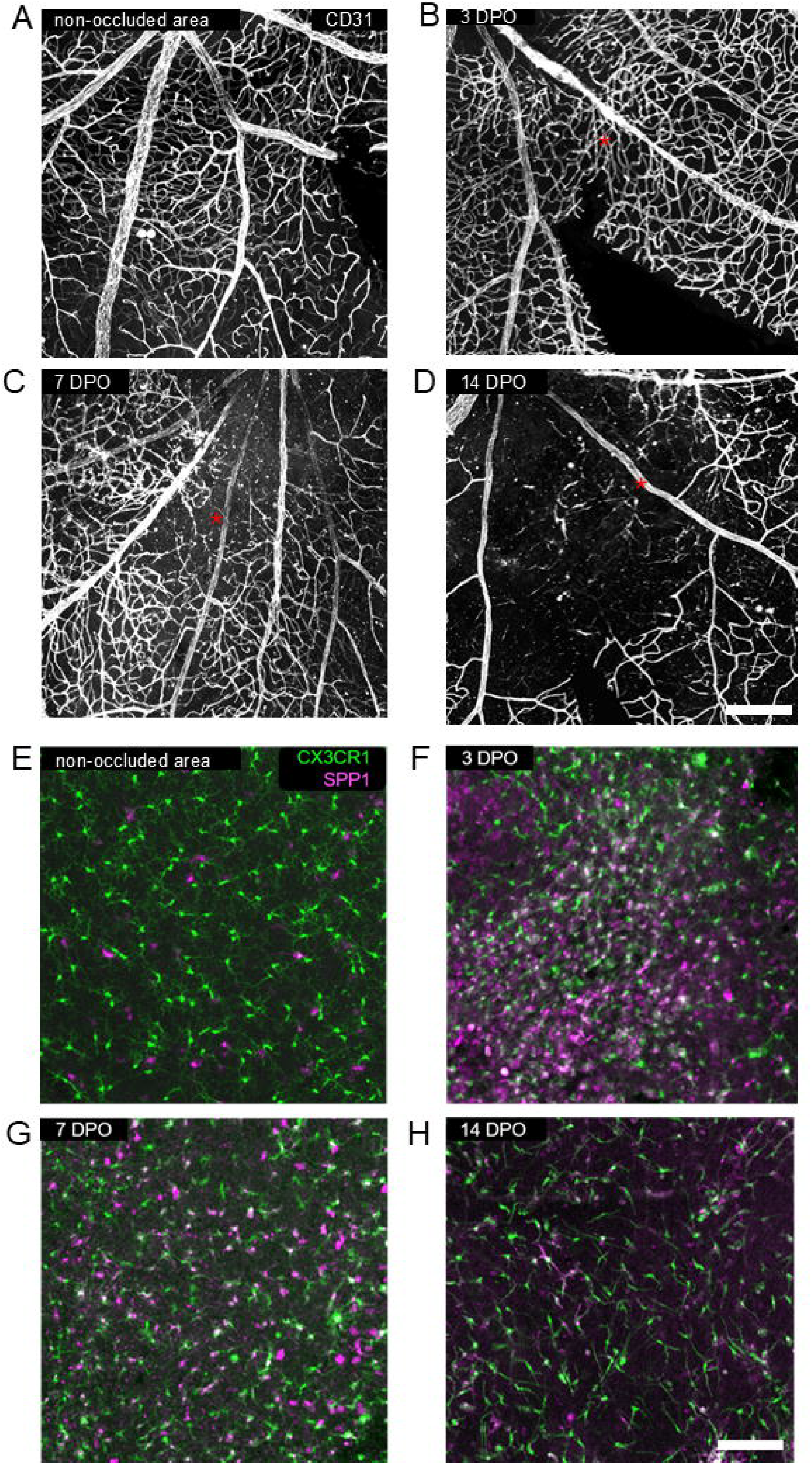
Vascular degradation and microglial activation in ischemic regions following BRVO. (A–D) Confocal maximum intensity projections of the superficial, intermediate and deep vascular plexuses, labeled with the endothelial marker CD31 from the non-ischemic (A) or ischemic regions at the indicated time points after BRVO. Red asterisks indicated photocoagulation site. (E–H) Confocal average intensity projections of the GCL and IPL labeling microglia with the Cx3Cr1-GFP transgenic mouse (green) and SPP1 (magenta) at the indicated time points after BRVO. Scale bars = 60 µm.

To examine the inflammatory response following ischemic induction, we monitored microglial recruitment, over the two-week ischemic period using CX3CR1-GFP mice (Fig. 2E-H). We detected more microglia in the occluded areas as early as 3 DPO, with maximal accumulation within the occluded areas at 7 DPO and substantial resolution by 14 DPO. Activated microglia were observed demonstrating characteristic amoeboid morphology, with enlarged cell bodies and simplified arbors.^33^ To identify reactive microglia, we evaluated SPP1 expression by immunostaining at 3, 7, and 14 DPO. At 3 DPO, SPP1 showed strong expression in microglia within the occluded areas (Fig. 2F), suggesting acquisition of a disease-associated inflammatory phenotype.^34^ However, in the non-occluded regions of BRVO eyes, we did not detect SPP1 expression in microglia, acquisition of an ameboid morphology, or disruption of their characteristic tiling pattern (Fig. 2E). Similarly, Sham-operated animals showed no significant microglial accumulation, morphological activation, or SPP1 upregulation, consistent with the findings in healthy eyes (Data not shown).

### 3.2 RGCs die continuously over two weeks after BRVO

We evaluated the dynamics of RGC loss after BRVO by immunostaining for the pan-RGC marker RBPMS at 3 time points following occlusion and then quantifying density in the occluded and non-occluded regions to measure a survival ratio. Within the BRVO affected region, RGCs died progressively over the 2-week time course (Fig. 3). Cumulative RGC losses were 32.0% ± 12.1% at 3 DPO, 60.9% ± 17.4% at 7 DPO, and 81.8% ± 5.8% at 14 DPO. RGC loss was significant from 3 DPO to 7 DPO (p = 0.0001), and from 7 DPO to 14 DPO (p = 0.0010) (Fig. 3F, left). These findings show that RGC death is not limited to the early phase after BRVO. Meanwhile, to evaluate the effect of photocoagulation on other retinal layers, we assessed TRT at 3, 7, and 14 DPO by Dapi staining (Fig. 3E). However, no statistically significant differences were observed among the groups (Fig. 3F, right), suggesting neuronal death was mostly limited to RGCs.

**Figure 3.**
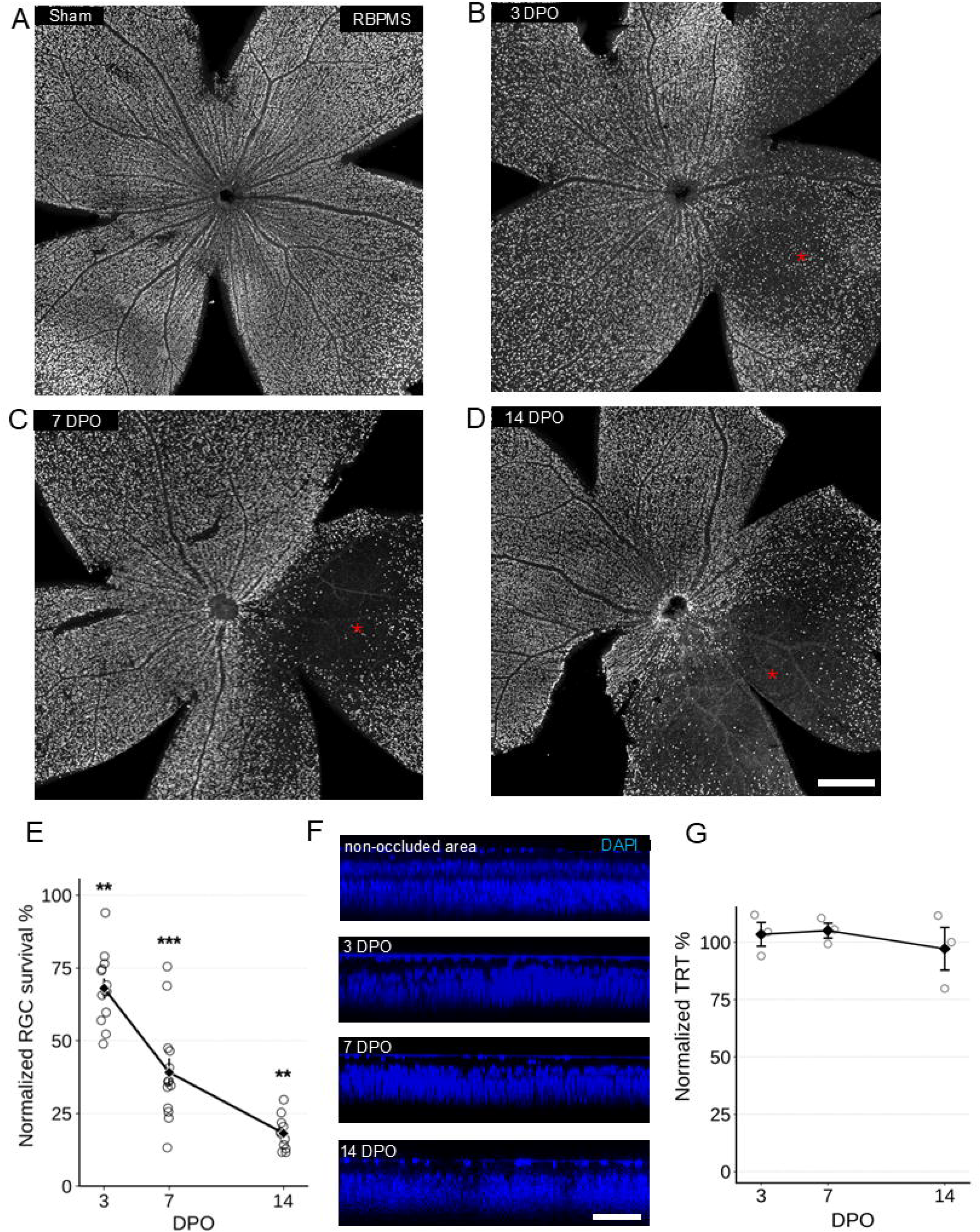
RGC death following BRVO. Confocal maximum intensity projections of retinal wholemounts immunostained for the pan-RGC marker RBPMS in sham-operated animals (A) or at 3, 7, and 14 DPO (B–D respectively). Scale bar = 300 µm. Red asterisks indicate the site of laser photocoagulation. (E) Quantification of RGC survival expressed as the ratio of RBPMS^+^ cell density in the occluded areas to the non-occluded areas of the same retina (3 DPO: n = 13; 7 DPO: n = 13; 14 DPO: n = 11); each circle represents one eye and the data are shown as mean ± SD. Significance was assessed by Welch’s t-test against sham controls at each timepoint (* p < 0.05, ** p < 0.01, *** p < 0.001). (F) Orthogonal views of DAPI-stained retinal flatmounts showing the non-occluded areas and the occluded areas at 3, 7, and 14 DPO after BRVO (n = 3 per timepoint). Scale bar = 80 µm. (G) Quantification of TRT measured from orthogonal DAPI images, expressed as the ratio of TRT in the ischemic region to the non-ischemic region of the same retina (n = 3 per timepoint).

We confirmed that light exposure alone did not cause measurable RGC loss using a sham group that received the same light exposure without Rose Bengal injection. At 2 weeks after sham, RBPMS^+^ cell density was not statistically different from untreated retinas (4295.4 ± 219.4 vs. 4491.4 ± 242.4 cells/mm^2^, respectively, Welch’s t-test, p = 0.363). Therefore, cell loss in ischemic retinas was caused by vascular occlusion rather than light exposure (Supplementary Figure S1).

### 3.3 αRGCs, F-RGCs and OODSGCs, die rapidly and near-completely

To examine if RGC types are differentially susceptible to BRVO we evaluated the survival of 4 RGC types: α-RGCs, OODSGCs, F-RGCs and ipRGCs (Fig. 4). At 3 DPO, SPP1^+^ αRGCs showed marked early loss following BRVO, with a 74.9% ± 5.8% loss over the first 3 days (p = 0.016; d = 5.41) (Fig. 4A). CART^+^ OODSGCs in the occluded versus non-occluded areas had declined by 78.8% ± 7.2% (p < 0.001; d = 10.71). In contrast, FOXP2^+^ F-RGCs were less affected during the first 3 days, with only a 55.9% ± 6.3% loss (p = 0.004; d = 8.23). However, by 7 DPO, FOXP2^+^ RGCs had decreased sharply by 82.6% ± 9.4% (p = 0.003; d = 7.61), and SPP1^+^ RGCs by 82.3% ± 5.6% (p < 0.001; d = 9.53) (Fig. 4B, E), and CART^+^ RGCs by 83.5% ± 7.2% (p = 0.030). At 14 DPO, CART^+^, SPP1^+^, and FOXP2^+^ RGC populations were all severely depleted. SPP1^+^ RGCs showed a 93.4% ± 3.5% loss, and FOXP2^+^ RGCs a 91.6% ± 5.4% loss, and CART^+^ RGCs an 86.0% ± 7.4% loss.

**Figure 4.**
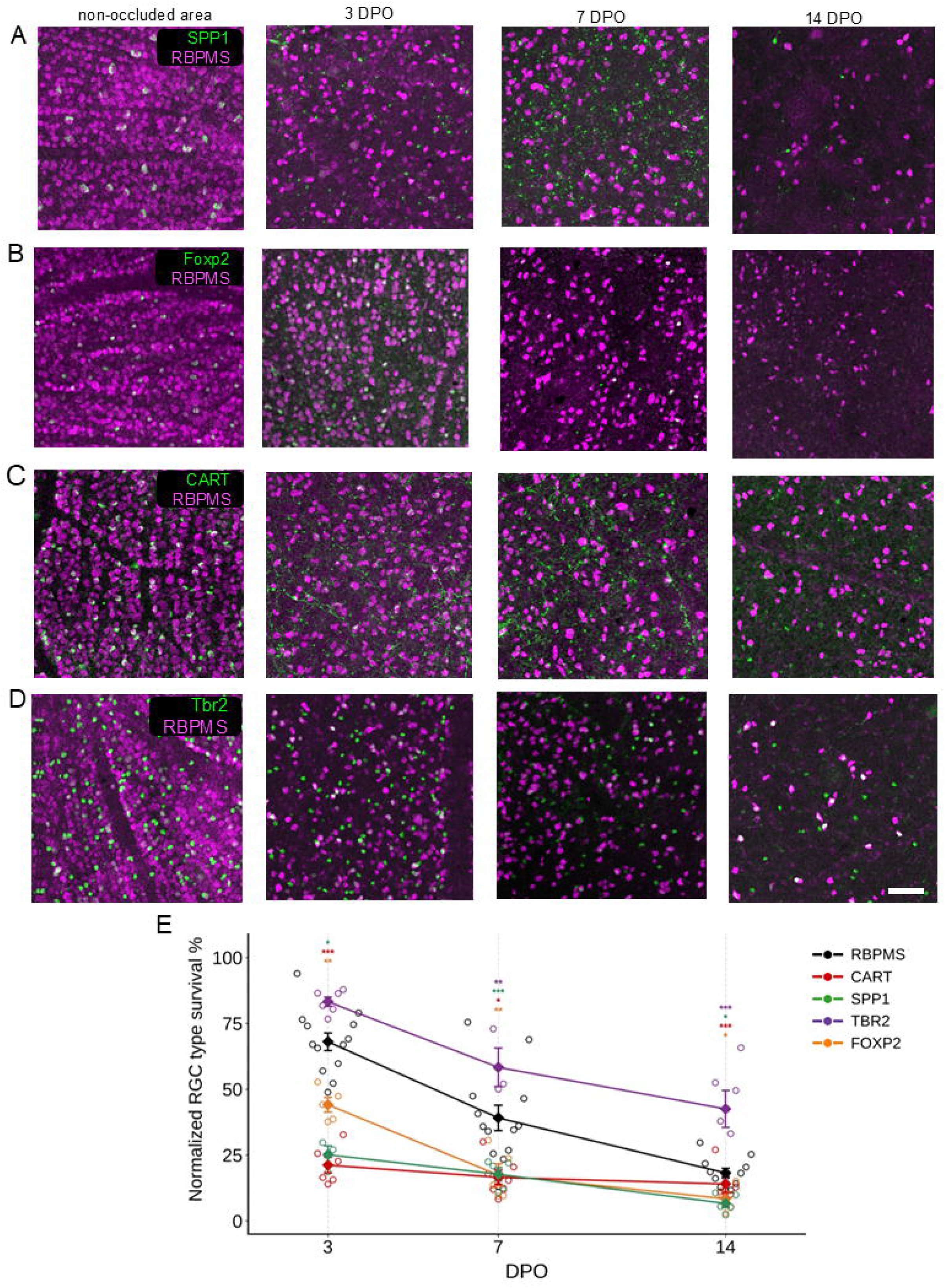
Subtype-specific vulnerability of RGCs after BRVO. Subtype-specific vulnerability of RGCs after BRVO. (A–D) Confocal maximum intensity projections of retinal wholemounts immunostained for the indicated RGC subtype marker (green) and the pan-RGC marker RBPMS (magenta) from the non-ischemic (A) or ischemic regions at 3, 7, and 14 DPO after BRVO. Scale bar = 100 µm. (E) Quantification of RGC subtype survival expressed as the ratio of marker^+^ RGC density in the occluded area to the non-occluded area of the same retina: RBPMS (black; 3 DPO: n = 13; 7 DPO: n = 13; 14 DPO: n = 11), SPP1^+^ αRGCs (green; 3 DPO: n = 3; 7 DPO: n = 5; 14 DPO: n = 5), CART^+^ OODSGCs (red; 3 DPO: n = 6; 7 DPO: n = 7; 14 DPO: n = 5), TBR2^+^ RGCs (purple; 3 DPO: n = 6; 7 DPO: n = 3; 14 DPO: n = 6), and FOXP2^+^ F-RGCs (orange; 3 DPO: n = 5; 7 DPO: n = 5; 14 DPO: n = 4). Each circle represents one eye and is color-coded to its respective group; the data are shown as mean ± SD. Significance was assessed by Welch’s t-test against sham controls at each timepoint (* p < 0.05, ** p < 0.01, *** p < 0.001).

### 3.4 ipRGCs are resilient to BRVO

At 3 DPO, TBR2^+^ RGCs showed only a 16.8% ± 4.4% loss (p = 0.180; d = 2.04), which did not reach statistical significance compared to sham and was significantly less than the overall RGC population loss at the same timepoint (32.0% ± 12.1%; p = 0.001). At 7 DPO, TBR2^+^ RGCs showed a 41.7% ± 12.7% loss (p = 0.004; d = 4.86), compared to 60.9% ± 17.4% loss in RBPMS^+^ RGCs, though this difference did not reach significance at this timepoint (p = 0.094). By 14 DPO, TBR2^+^ RGCs showed a 57.5% ± 17.2% loss — significantly less than the overall RGC population loss (81.8% ± 5.8%; p = 0.017) or any other RGC subtype examined (SPP1^+^ (p = 0.003), FOXP2^+^ (p = 0.003), and CART^+^ RGCs (p = 0.008)) (Fig. 4D, E).

## Discussion

Clinical studies indicate that hemodynamic changes in BRVO can affect the retina in a gradient of hypoperfusion. Previous clinical studies reported decreases in blood flow and vessel density in both occluded and non-occluded retinal regions in BRVO eyes.^35,45^ Similarly, in our study, we observed a significant decrease in perfusion and vascular structure in the occluded region 2 days after occlusion, while the non-occluded region also showed vascular changes, but with less severity. Since we performed *in vivo* angiography and perfusion measurements at an acute time point, we do not know how local perfusion changed over the longer term. However, the marked vascular destruction that developed over 14 days should induce a substantial decrease in perfusion of the occluded region.

It should be noted that the mismatch between capillary density measured by OCTA and histology may be explained by differences in the methods. OCTA measures blood flow or perfusion, whereas histology shows vessels. Therefore, the decrease in perfusion may occur before visible loss of vascular structure. Several mechanisms have been proposed to explain the vascular and hemodynamic damage in both occluded and non-occluded retinal regions including leukocyte adhesion, effects of diffusible mediators such as VEGF, pericyte loss, and changes in vascular tone and autoregulation.^10, 36^ In addition, off-target activation of Rose Bengal within the capillaries following photocoagulation could be another potential contributor to the vascular destruction observed in our experiment. However, we would expect more widespread capillary damage, particularly in the deeper retinal layers, if this were the case.

Vascular instability can cause fluid leakage, leading to retinal edema and changes in retinal layer thickness.^37^ Similarly, in our study, total retinal thickness increased in both occluded and non-occluded regions, along with sublayer changes (Fig. 3E). How ever the results were not statistically significant, likely because our experiments were underpowered for the low resolution measurements.

Microglial migration and activation have been reported in laser-induced CNV, diabetic retinopathy, and oxygen-induced retinopathy (OIR).^38–40^ In OIR, which involves ischemia across the retina, Spp1 is enriched in activated microglial clusters and is associated with increased accumulation of retinal myeloid cells around neovascular areas.^41^ Also, Spp1 expression is regulated by hypoxia-inducible factor 1 (HIF-1a) and NF-κB signaling pathways and contributing to neovascularization.^34^ In CNV models, SPP1 is among the most highly expressed genes by retinal microglia, and upregulation was associated with a disease-associated amoeboid morphology. However, intravitreal anti-SPP1 treatment increased CNV lesion size.^42^ In contrast, a separate study reported that systemic anti-SPP1 antibody treatment reduced VEGF mRNA expression and macrophage invasion in the choroidal tissue, resulting in smaller CNV lesions.^43^

Overall, SPP1 has been reported to increase in retinal and choroidal ischemic injuries and may contribute to disease progression through effects on vascular proliferation, endothelial cell migration, and CD44 signaling in the retina.^44^ In our experiment, we observed microglial accumulation as early as 3 DPO in the occluded region, together with strong local SPP1 expression. In contrast, the non-occluded region maintained the native microglial arrangement, suggesting that the inflammatory response was mainly localized to the occluded region. At later time points, microglial density appeared to decrease along with SPP1 expression.

We evaluated RGC subtype survival in a mouse model of BRVO at 3, 7, and 14 DPO. The total number of RGCs dropped quickly and continued to decrease over the two weeks following ischemia. Subtype analysis showed that ipRGCs had the highest survival in the ischemic retina at all three time points. In contrast, αRGCs and F RGCs were the most susceptible subtypes in BRVO, while OODSGCs showed slightly better survival than these two groups.

RGCs are physiologically and molecularly heterogeneous, and they demonstrate different responses to degeneration. In models of axonal damage, glaucoma, and excitotoxicity, both ipRGCs and αRGCs have been included among the relatively resistant populations. Furthermore, SPP1 overexpression combined with IGF-1 or BDNF can promote mTOR-dependent axon regeneration, particularly in αRGCs, following optic nerve crush.^45^ Increased SPP1 levels in the retina have also been reported to have protective effects on RGCs in models of glaucoma, aging, and excitotoxicity.^46,47^ We found that ipRGCs were the most resistant subtype to ischemic injury. Surprisingly, αRGCs showed rapid cell death following ischemia and almost completely disappeared from the site of BRVO by 14 DPO. These findings suggest that RGC type survival differences depend on the type or etiology of degeneration. Although the mechanism underlying the high vulnerability of αRGCs during ischemia is not clear, it has been shown that αRGCs are reliant on oxidative phosphorylation to produce ATP and the RGC type most sensitive to mitochondrial disruption.^22^ This suggests that αRGCs may be more prone to energetic collapse during ischemia.

Upstream pathways may also contribute to αRGC vulnerability. According to transcriptomic data reported by Tran et al., αRGCs show relatively high basal Hif-1a expression, which translate to a stronger HIF-1a response during retinal ischemia.^48^ Excessive HIF-1a activation under severe ischemic conditions can become harmful by activating pro-death transcriptional programs rather than directly causing metabolic failure since HIF-1a overexpression increases neuronal death during oxygen-glucose deprivation, while HIF-1a inhibition or knockdown reduces neuronal death.^49^ Blocking HIF-1a activity can also protect cortical neurons from delayed hypoxic death in culture, while brain-specific Hif-1a deletion reduces neuronal apoptosis after acute hypoxic-ischemic injury and reduced the expression of pro-apoptotic genes.^50,51^ Future experiments focusing on the dynamics of transcription factors and pathways, including HIF-1a, across different RGC subtypes may help explain the high susceptibility of αRGCs.

## Supporting information

Sham group

## Acknowledgements

This research was supported by Research to Prevent Blindness (Career Development Award, PRW), NIH Grants EY032908, EY032908 and EY036111 (PRW), the Richard and Isabel Goldstein Family Foundation (PRW), the Jeffrey T. Fort Innovation Fund (RSA), and the Siteman Retina Research Fund (RSA). John F. Hardesty, MD Department of Ophthalmology and Visual Sciences is supported by NIH P30 EY002687 Core Grant and an unrestricted grant from Research to Prevent Blindness, Inc, New York, NY.

