## Supplementary material for "Differential Survival of Retinal Ganglion Cell Types Following Branch Retinal Vein Occlusion": Sham group

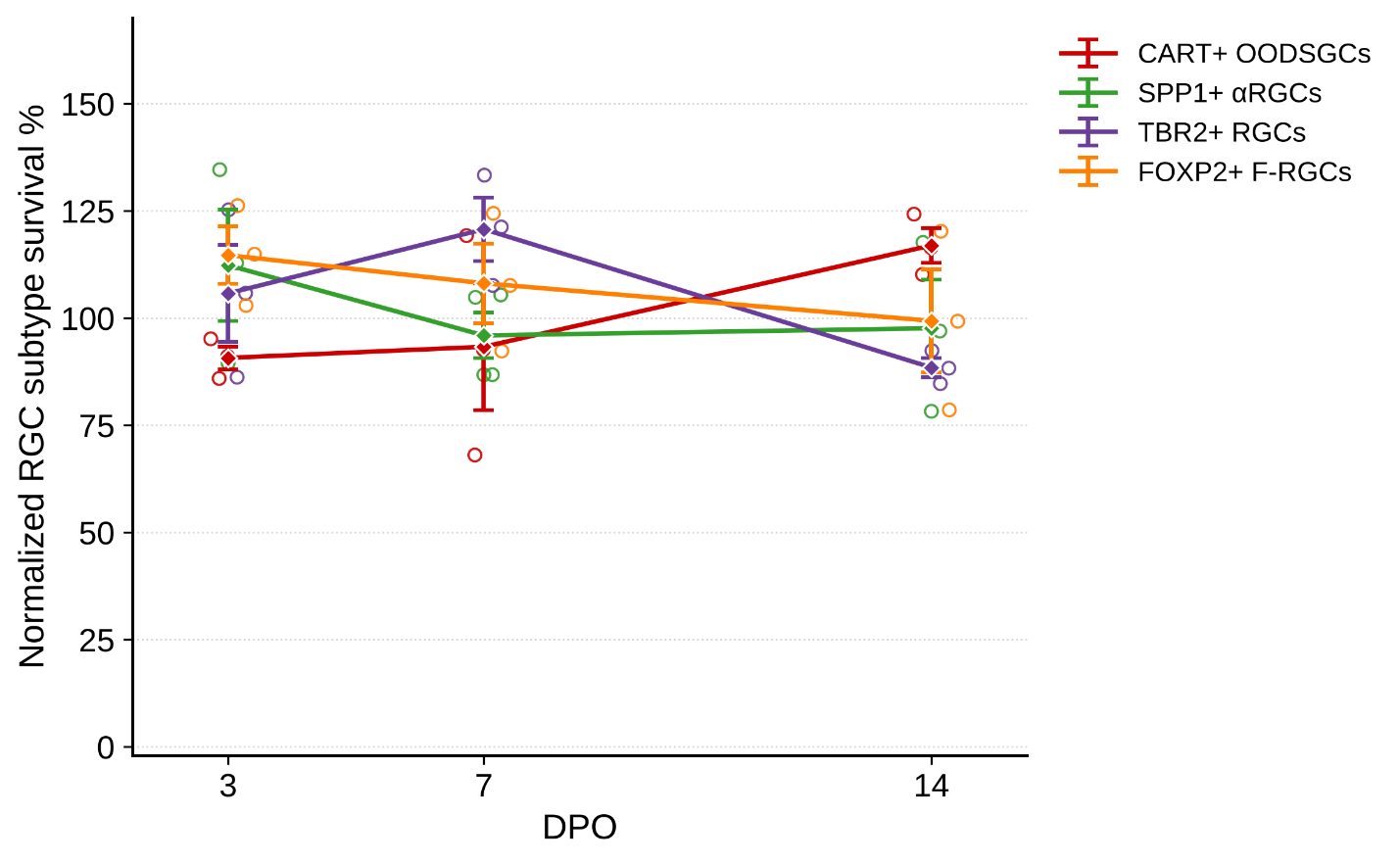
**Supplementary Figure 1. RGC subtype survival ratios in sham-operated animals.** Normalized survival of RGC subtypes in sham mice that received laser photocoagulation without Rose Bengal injection, at 3, 7, and 14 days post-operation (DPO). The ischemic-to-healthy region density ratio (I/H) is expressed as a percentage, with 100% indicating no difference between regions. Open circles represent individual animals; diamonds show mean ± SEM. The dashed line at 100% denotes equal density in both regions. No significant deviation from baseline was observed for any subtype at any timepoint. SPP1⁺ αRGCs: n = 3, 4, 3; TBR2⁺ RGCs: n = 3, 3, 3; FOXP2⁺ F-RGCs: n = 3, 3, 3; CART⁺ OODSGCs: n = 3, 3, 3 at 3, 7, and 14 DPO respectively.
